# β-Lapachone Regulates Mammalian Inositol Pyrophosphate Levels in an NQO1- and Oxygen-dependent Manner

**DOI:** 10.1101/2022.11.27.518098

**Authors:** Verena B. Eisenbeis, Danye Qiu, Lisa Strotmann, Guizhen Liu, Isabel Prucker, Kevin Ritter, Christoph Loenarz, Adolfo Saiardi, Henning J. Jessen

## Abstract

1

Inositol pyrophosphates (PP-InsPs) are energetic signalling molecules with important functions in mammals. As their biosynthesis depends on ATP concentration, PP-InsPs are tightly connected to cellular energy homeostasis. Consequently, an increasing number of studies involves PP-InsPs in metabolic disorders, such as type 2 diabetes, aspects of tumorigenesis, and hyperphosphatemia. Research conducted in yeast suggests that the PP-InsP pathway is activated in response to reactive oxygen species (ROS). However, the precise modulation of PP-InsPs during cellular ROS signalling is unknown. Here, we report how mammalian PP-InsP levels are changing during exposure to exogenous (H_2_O_2_) and endogenous ROS. Using capillary electrophoresis electrospray ionization mass spectrometry (CE-ESI-MS), we found that PP-InsP levels decrease upon exposure to oxidative stressors in HCT116 cells. Application of quinone drugs, particularly β-lapachone (β-lap), under normoxic and hypoxic conditions enabled us to produce ROS *in cellulo* and to show that β-lap treatment caused PP-InsP changes that are oxygen dependent. Experiments in MDA-MB-231 breast cancer cells deficient of NAD(P)H:quinone oxidoreductase-1 (NQO1) demonstrated that β-lap requires NQO1-bioactivation to regulate the cellular metabolism of PP-InsPs. Critically, significant reductions in cellular ATP concentrations were not directly mirrored in reduced PP-InsP levels as shown in NQO1-deficient MDA-MB-231 cells treated with β-lap. The data presented here unveil new aspects of β-lap pharmacology and its impact on PP-InsP levels. Our identification of different quinone drugs as modulators of PP-InsP synthesis will allow to better appreciate their overall impact on cellular function.

**Significance Statement:** Inositol pyrophosphates (PP-InsPs) are messenger molecules regulating various functions in mammals. They are associated with the oxidative stress response, but the underlying mechanism is unclear. We investigate PP-InsP signalling in mammalian cells subjected to reactive oxygen species (ROS). Applying the quinone β-lapachone (β-lap) generated intracellular ROS resulting in decreased PP-InsP levels. This indicates a key role of PP-InsPs in cellular signalling under oxidative stress. Moreover, β-lap mediated PP-InsP signalling required oxygen and the enzyme NAD(P)H:quinone oxidoreductase-1 (NQO1). Since quinone drugs are cytotoxic, our data provide a basis for further investigations into the role of PP-InsPs during quinone-dependent chemotherapies. This is of special relevance since a phase II clinical trial demonstrated β-lap efficacy in a combination chemotherapy against pancreatic cancer.

## 3 Introduction

*myo*-Inositol pyrophosphates (PP-InsPs hereafter) are intracellular messengers implicated in a wide range of physiological processes in eukaryotes. Particularly, they have been referred to as “metabolic messengers” as their concentration is bound to ATP levels.^[1–4]^ They are composed of phosphate esters and either one or two pyrophosphate groups attached to the six-carbon *myo*-inositol ring resulting in PP-InsP_5_ or (PP)_2_-InsP_4_, respectively.^[1]^ Apart from regular protein binding,^[2]^ PP-InsPs are also capable of nonenzymatic protein pyrophosphorylation.^[5, 6]^ Mammalian PP-InsP synthesis mainly starts from the inositol phosphate InsP_6_ and is achieved by two different classes of enzymes: the inositol hexakisphosphate kinases (IP6Ks) and the diphosphoinositol-pentakisphosphate kinases (PPIP5Ks, **Figure 1 A**).^[1, 4]^ Among the biosynthesized PP-InsPs, 5-PP-InsP_5_ is the most abundant isomer inside mammalian cells while 1-PP-InsP_5_ and 1,5-(PP)_2_-IP_4_ occur in comparably low concentrations.^[1]^ However, very recent data suggests that large fluctuations in concentrations of the distinct isomers can occur in mammalian tissues and that additional isomers are also present.^[7]^

**Figure 1.**
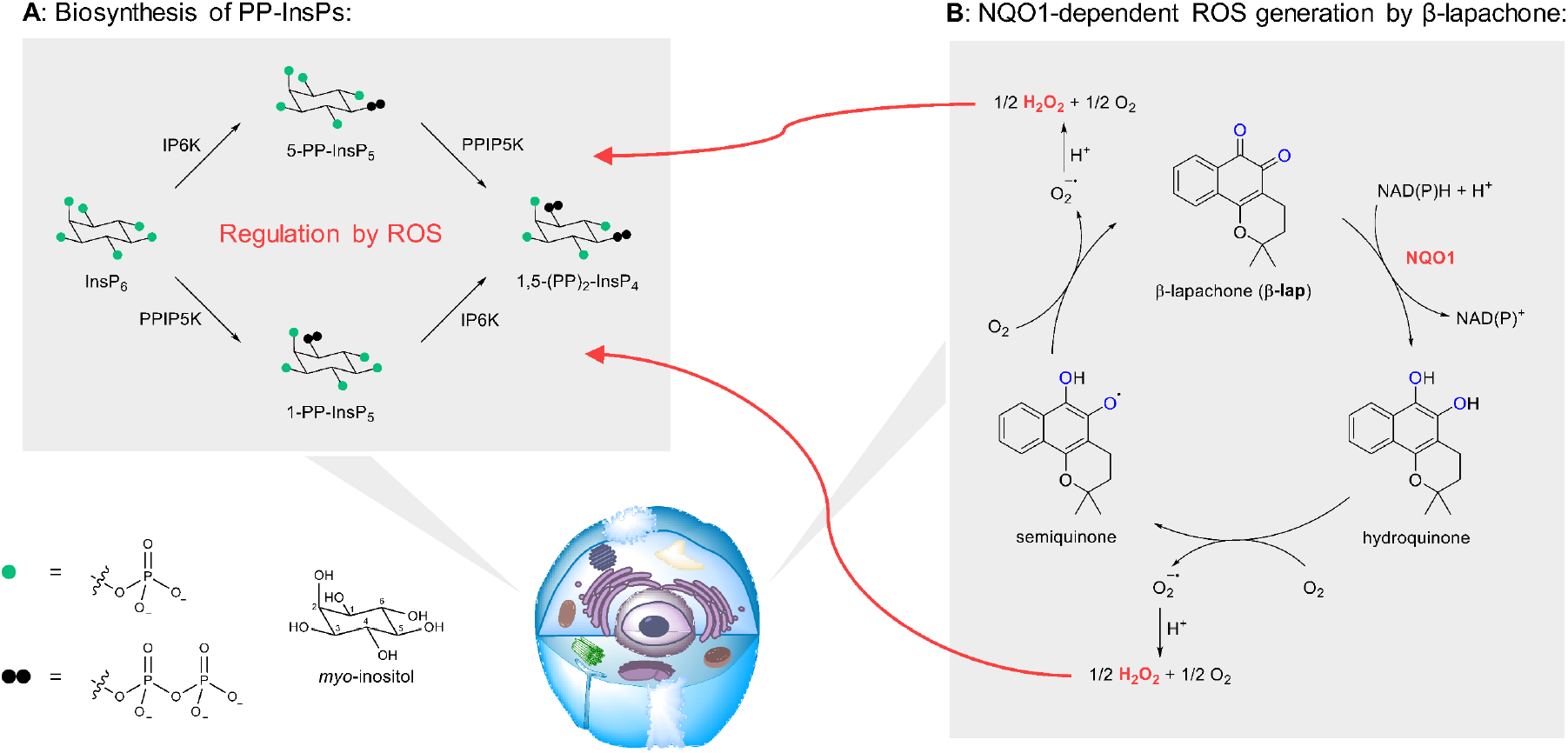
Exploring the influence of ROS on mammalian PP-InsP levels. (***A**) Biosynthesis of PP-InsPs from InsP_6_ via IP6Ks and PPIP5Ks. (**B**) Proposed mechanism of action of β-lap*. β-Lap redox cycles in NQO1 expressing cells between hydroquinone and semiquinone forms thereby generating cytotoxic ROS. Spontaneous hydroquinone and semiquinone oxidation occurs in the presence of oxygen.

To date, a variety of studies have connected PP-InsPs and their kinases with diseases, such as type 2 diabetes,^[8, 9]^ carcinogenesis,^[10, 11]^ and hyperphosphatemia.^[12]^ New therapeutic strategies targeting the PP-InsP pathway are therefore evolving.^[12]^ Mammals express three different IP6K paralogs: IP6K1/2/3.^[2]^ Among those, IP6K2 has attracted significant attention since several experiments demonstrated that this isoform sensitizes mammalian cells to apoptosis caused by stressors such as reactive oxygen species (ROS), *γ*-irradiation, and cisplatin.^[13, 14]^ Moreover, it was found that ROS decreased PP-InsP_5_, as well as (PP)_2_-InsP_4_ levels in yeast cells suggesting that PP-InsPs are directly involved in the yeast oxidative stress response.^[15]^

As massive oxidative stress is lethal to the cell, chemotherapeutic agents that selectively release ROS in cancers have been identified. The naphthoquinone β-lapachone (β-lap) is an antitumor drug with activity against solid tumours evaluated in several phase I clinical trials.^[16, 17]^ Furthermore, it showed efficacy against pancreatic cancer in a phase II study when applied in combination with gemcitabine.^[18]^ β-Lap is capable of releasing ROS in cells expressing the two-electron reductase NAD(P)H:quinone oxidoreductase 1 (NQO1, EC 1.6.5.2).^[19]^ According to the mechanism proposed by Pink *et al*., NQO1 reduces the drug to an unstable hydroquinone thereby consuming one molecule NAD(P)H per reaction. The hydroquinone autoxidizes through a semiquinone intermediate to the parent compound. This spontaneous oxidation produces ROS, closes the futile cycle, and serves as starting point for the next round (**Figure 1 B**).^[20]^ Using MCF7:WS8 cell extracts as NQO1 source it was demonstrated, that one mole of β-lap was capable of oxidizing 50-60 mole NADH in 3 min.^[20]^ As various human solid tumors express significantly higher NQO1 levels than normal tissues,^[19]^ β-lap enables selective killing of cancer cells.

β-Lap mediated cellular ROS generation has been extensively studied^[21–25]^ and various downstream targets have been described. These include hyperactivation of poly(ADP-ribose)polymerase-1 in an NQO1-dependent manner,^[23]^ which leads to dramatic decrease of intracellular ATP.^[23, 26, 27]^ A recently developed prodrug-based delivery technique that improves tumor targeting demonstrates that systemic toxicity can be significantly reduced, rendering *o*-quinones a candidate compound family for targeted therapeutics.^[28]^

In this work, we explore the levels of mammalian PP-InsPs in response to oxidative stress, particularly brought about by quinone reagents, such as β-lap, in an NQO1-dependent manner under normoxic and hypoxic conditions (overview shown in **Figure 1**). We therefore add another layer to our understanding of *o*-quinone action on cells. Application of highly sensitive capillary electrophoresis electrospray ionization mass spectrometry (CE-ESI-MS) allowed the absolute quantitation of PP-InsP levels in two different mammalian cell lines.^[29]^ These cell lines either expressed (HCT116^UCL^ cells) or did not express NQO1 (MDA-MB-231 cells). We demonstrate the importance of NQO1 for the modulation of PP-InsP levels under hypoxic and normoxic conditions. With the emergence of targeted *o*-quinones as cytotoxic drugs,^[28]^ a deeper understanding of their impact on cell signalling molecules is highly warranted and provided in this study.

## 4 Results and Discussion

### H_2_O_2_ and bioactivated β-lapachone reduce PP-InsP levels in HCT116^UCL^ cells

To evaluate the impact of exogenous ROS on mammalian PP-InsP levels, the human colon cancer cell line HCT116^UCL^ was used.^[30]^ HCT116^UCL^ cells maintain higher 5-PP-InsP_5_ as well as 1,5-(PP)_2_-InsP_4_ levels than other mammalian cell lines^[31]^ and are consequently ideally suited to study changes in PP-InsP concentrations. These analytes can be readily detected using CE-ESI-MS with very high resolution.^[7, 32, 33]^

Initially, HCT116^UCL^ cells were oxidatively stressed *via* exposure to 0.1 mM or 1.0 mM H_2_O_2_ for 30 min. Then, the cellular metabolism was directly quenched with perchloric acid and PP-InPs and InsP_6_ were enriched with TiO_2_ beads.^[34]^ The enriched extracts were spiked with stable isotope labelled (SIL) [^13^C]-PP-InsP and [^13^C]-InsPe_6_^[35, 36]^ internal standards and analyzed by CE-ESI-MS.^[37]^ Detection of 1,5-(PP)_2_-InsP_4_, 5-PP-InsP_5_, and InsP_6_ was feasible, whereas 1-PP-InsP_5_ levels were below the detection limit in several samples and therefore not further considered for quantitation. (**Figure 2 A**). Normalization by protein content revealed decreased 5-PP-InsP_5_ and 1,5-(PP)_2_-InsP_4_ levels after treatment with 1.0 mM H_2_O_2_. PP-InsP levels in HCT116^UCL^ cells subjected to 0.1 mM H_2_O_2_ as well as InsP6 levels in cells treated with either 0.1 mM or 1.0 mM H_2_O_2_ were not affected (**Figure 2 B1** and **B2**).

**Figure 2.**
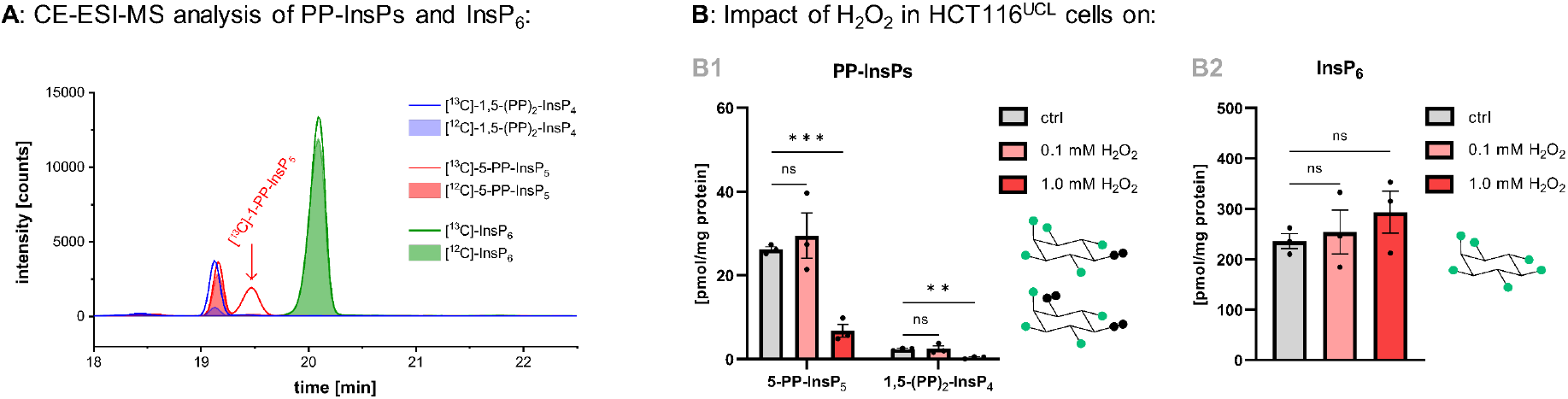
Exogenous ROS (H_2_O_2_) decrease mammalian PP-InsP levels. (***A**) Electropherogram of 1,5-(PP)_2_-InsP_4_, 5-PP-InsP_5_, and InsP_6_ (filled areas) in untreated HCT116^UCL^ cells and stable isotope labelled [^13^C]-1,5-(PP)_2_-InsP_4_, [^13^C]-5-PP-InsP_5_, [^13^C]-1-PP-InsP_5_, and [^13^C]-InsP_6_ internal standards (unfilled areas)*. 1-PP-InsP_5_ in HCT116^UCL^ control cells was below the detection limit in several samples and therefore not considered for quantitation. (***B**) PP-InsP and InsP_6_ levels in HCT116^UCL^ control cells and HCT116^UCL^ cells treated with 0.1 mM or 1.0 mM H_2_O_2_ for 30 min*. Data are means ±SEM from three replicates. Statistical analyses to compare treated cells with control cells were performed using an unpaired two-tailed student’s t-test (\*\**P* ≤ 0.01, \*\*\**P* ≤ 0.001). ns: not significant. ctrl: control.

These results are in agreement with earlier studies in yeast revealing reduced PP-InsP levels after incubation with 1 mM H_2_O_2_ for 20 min.^[15]^ However, Choi *et al*. described unaffected (PP)2-InsP4 levels in DDT_1_ MF_2_ cells after treatment with 0.15 mM or 0.3 mM H_2_O_2_ for 30 min.^[38]^ We conclude that extracellular H_2_O_2_ concentrations in the millimolar range are required to significantly lower PP-InsP levels in mammalian cell lines, particularly HCT116^UCL^. Analysis of methanol-extracted ATP from co-cultured cells revealed unaffected ATP levels in cells incubated with 0.1 mM H_2_O_2_. In contrast, ATP levels in HCT116^UCL^ cells treated with 1.0 mM H_2_O_2_ were reduced (ca. 3-fold) confirming reported cellular ATP losses after excessive exposure to oxidative stress (**Figure S1**).^[39]^ The oxidative stress-dependent reduction of PP-InsP levels as a potential response to reduced ATP levels might be the result of an interplay between PP-InsP signalling and cellular energy dynamics.^[2, 3]^ However, it remains unclear if ATP reduction or other effects of ROS are critically required for PP-InsP reduction upon H_2_O_2_ exposure.

Therefore, to get a deeper understanding of ROS-dependent PP-InsP regulation, we next investigated whether H_2_O_2_ mediated PP-InsP changes had been caused by intracellular ROS. The bioactivatable quinone β-lap was chosen as an intracellular generator of oxidative stress. HCT116 cells are synthesizing functioning NQO1^[40, 41]^ and endogenous ROS production in this cell line can consequently be achieved by β-lap treatment. To test whether PP-InsP levels were altered upon exposure, HCT116^UCL^ cells were subjected to 2.5 μM, 5 μM, or 10 μM β-lap for 2 h. Cellular metabolism was then quenched and 5-PP-InsP_5_, 1,5-(PP)_2_-InsP_4_, and InsP_6_ levels were analyzed by CE-ESI-MS. β-Lap dose-dependently decreased PP-InsP levels while InsP_6_ levels remained stable (**Figure 3 A1-A3**). Treatment with 10 μM of the naphthoquinone for 2 h almost completely depleted cellular PP-InsP levels. In addition, ATP from cells grown on parallel dishes and subjected to 5 μM or 10 μM of β-lap was analysed and β-lap mediated depletion of ATP levels was confirmed (**Figure S2 A**).^[23, 26, 27]^ The results obtained from assaying PP-InsPs, InsP_6_, and ATP levels in β-lap treated HCT116^UCL^ cells supported the idea that mammalian PP-InsPs are responding to intracellularly generated oxidative stress, whereas InsP_6_ remains unaffected.

**Figure 3.**
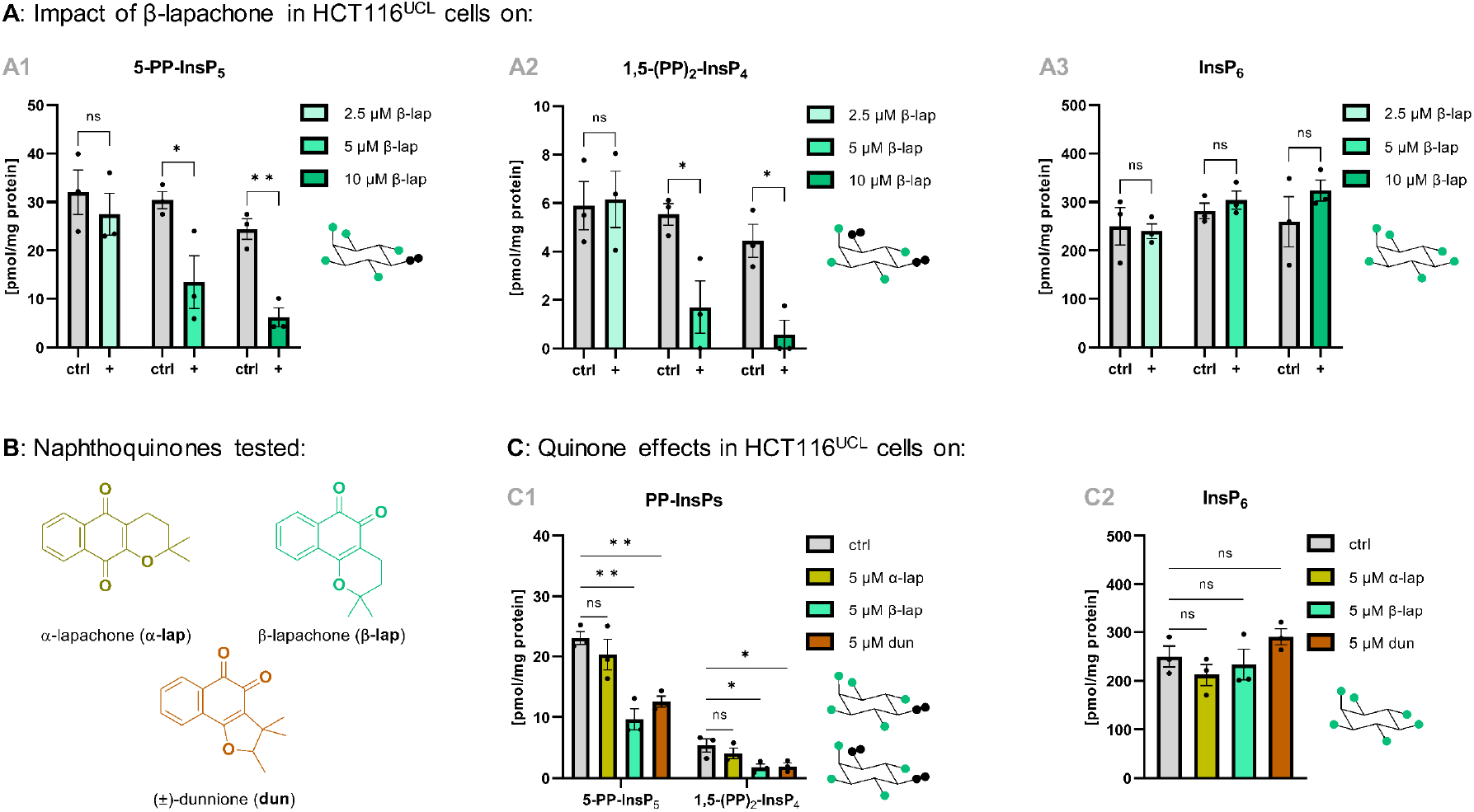
Mammalian PP-InsP levels fall in response to naphthoquinone-generated endogenous ROS. *PP-InsP and InsP_6_ levels in HCT116^UCL^ control cells and HCT116^UCL^ cells treated with 2.5 μM, 5 μM, or 10 μM β-lap for 2 h. (**B**) Structures of the p-naphthoquinone α-lap and the o-naphthoquinones β-lap and dun used in this study. (**C**) PP-InsP and InsP_6_ levels in HCT116^UCL^ control cells and HCT116^UCL^ cells treated with 5 μM α-lap, β-lap, or dun for 2 h*. (**A**)+(**C**): Data are means ±SEM from three replicates. Statistical analyses to compare treated cells with control cells were performed using an unpaired two-tailed student’s t-test (\**P* ≤ 0.05; \*\**P* ≤ 0.01). ctrl: control. ns: not significant.

In addition to β-lap, other *o*-naphthoquinones, such as (±)-dunnione (dun, chemical structure shown in **Figure 3 B**), are substrates for NQO1 and are able to generate large quantities of ROS inside the cell.^[42]^ In contrast, the *p*-naphthoquinone α-lapachone (α-lap, chemical structure shown in **Figure 3 B**) is known to produce no or only comparably small amounts of intracellular ROS.^[43–45]^ Therefore, the impact of α-lap, β-lap, and dun on PP-InsP levels in HCT116^UCL^ cells was compared with their reported ability to generate cellular ROS. HCT116^UCL^ cells were incubated with 5 μM α-lap, β-lap, or dun for 2 h and PP-InsPs and InsP_6_ were extracted from the cells as described above. CE-ESI-MS analysis revealed that β-lap and dun were equally capable of reducing 5-PP-InsP5 (ca. 2-fold) and 1,5-(PP)_2_-InsP_8_ levels (ca. 3-fold) whereas α-lap had no significant impact on PP-InsP levels (**Figure 3 C1**). Neither the two different *o*-quinones nor the *p*-quinone altered InsP_6_ levels (**Figure 3 C2**). The naphthoquinone-dependent extent of reduction of PP-InsP levels was therefore in alignment with their ability to produce ROS inside the cell.^[42–45]^

The *in cellulo* studies with β-lap and its isomers indicated an oxidative stress-dependent effect of the quinone on PP-InsP levels. However, a potential direct binding of β-lap to mammalian IP6Ks resulting in reduced PP-InsP synthesis could not be ruled out. Hence, β-lap was screened against purified IP6K1 serving as representative paralog for the IP6K family. Interestingly, the *in vitro* experiments revealed a decrease in IP6K1 activity with an IC_50_ value of ca. 10 μM and a maximal inhibition of 70 % (**Figure S2 B**). These results demonstrate that β-lap could directly target IP6K1, rendering it one new inhibitor of IP6Ks of which only a handful exist.^[12, 46–48]^ However, the *in cellulo* reduction of PP-InsP levels by the quinone was already significant at a concentration as low as 5 μM (**Figure 3 A1** and **A2**) showing that, apart from minor contributions of direct inhibition, β-lap was likely affecting cellular PP-InsP levels through generation of ROS.

### Hypoxia reveals that β-lapachone acts through ROS production

To test our hypothesis, we investigated if PP-InsP levels are reduced as a result of oxidative stress triggered by futile cycling of β-lap. Since the autoxidation of β-lap is dependent on partial O_2_ pressure (p[O_2_]) (**Figure 1 B**), reduced p[O_2_] is expected to diminish β-lap mediated ROS production and consequently lower loss of PP-InsP levels. In initial experiments, basal 5-PP-InsP_5_, 1,5-(PP)_2_-InsP_4_, and InsP_6_ concentrations in HCT116^UCL^ cells cultured under hypoxia were determined. Cells were initially seeded and pre-incubated under normoxia. Then, the dishes were placed in a hypoxia chamber and incubated at 1 % O_2_ (referred to as hypoxic conditions) for different time intervals. Subsequently, cells were lysed followed by enrichment as well as CE-ESI-MS analysis of PP-InsPs and InsP_6_ (**Figure 4 A**). Analyses of cellular ATP extracted from parallel dishes were also performed.

**Figure 4.**
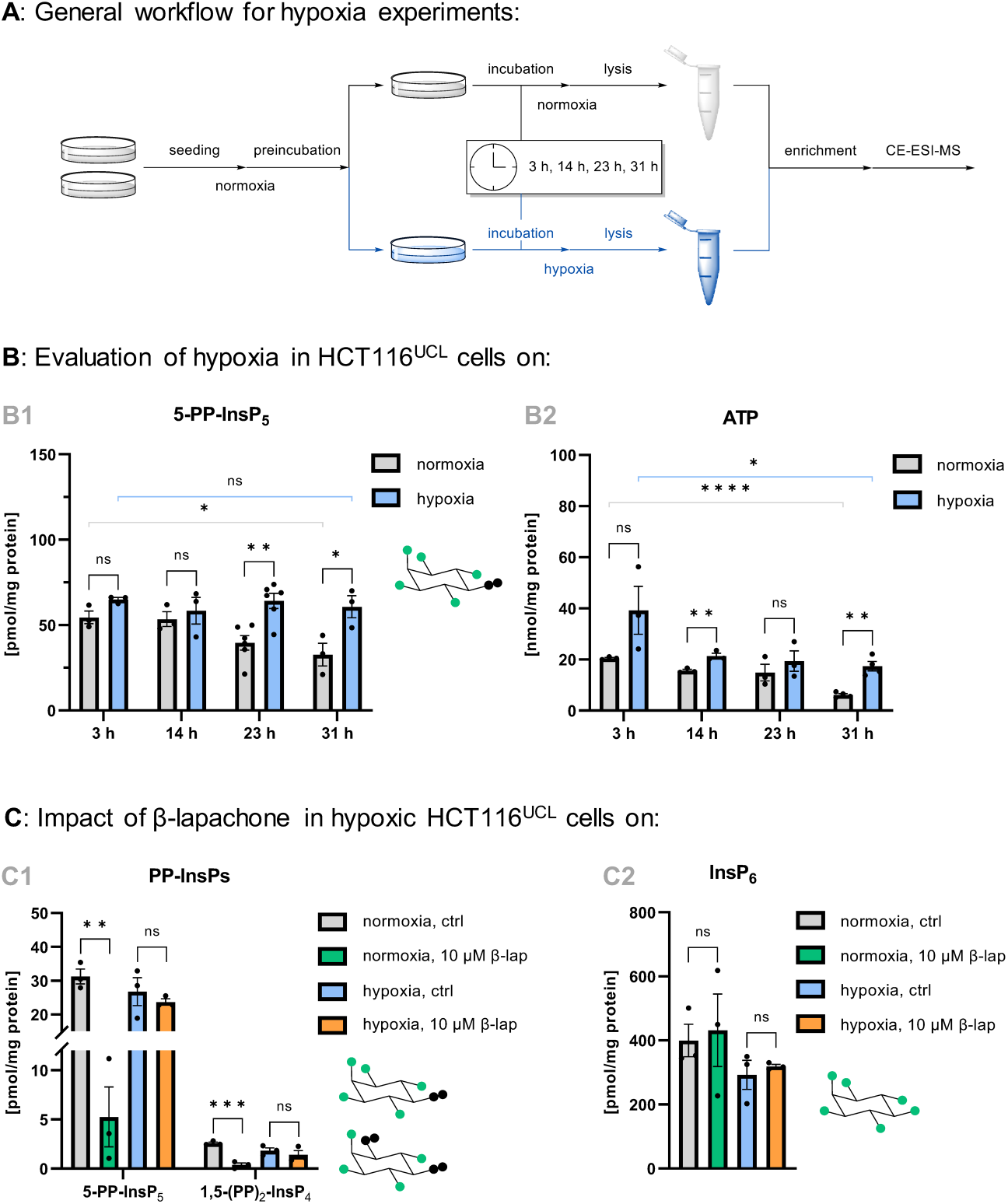
Oxygen is required for β-lapachone effects on PP-InsP levels. (***A**) Workflow for hypoxia experiments with subsequent enrichment of PP-InsPs and InsP_6_ from the cells followed by CE-ESI-MS. (**B**) 5-PP-InsP_5_ and ATP levels in HCT116^UCL^ cells grown for 3 h, 14 h, 23 h, and 31 h under normoxic and hypoxic conditions*. ATP was extracted from cells grown in parallel to those used for perchloric acid extration of PP-InsPs and InsP_6_. (***C**) PP-InsP and InP_6_ levels in normoxic and hypoxic control HCT116^UCL^ cells and normoxic and hypoxic HCT116^UCL^ cells treated with 10 μM β-lap for 2 h*. Preincubation time of cells under hypoxia prior to drug addition was 23 h. (**B**)+(**C**) Data are means ±SEM from three to six replicates. Statistical analyses to compare hypoxic with normoxic cells, cells cultured for 31 h under normoxia/hypoxia with cells cultured for 3 h under normoxia/hypoxia, and normoxic/hypoxic β-lap treated cells with normoxic/hypoxic control cells were performed using an unpaired twotailed student’s t-test (\**P* ≤ 0.05; \*\**P* ≤ 0.01; *\*\*\*P* ≤ 0.001; *\*\*\*\*P* ≤ 0.0001). ns: not significant. ctrl: control.

Interestingly, 5-PP-InsP_5_ levels in normoxic cells declined to about half of the initial value at the endpoint of this study (31 h). In contrast, 5-PP-InsP_5_ levels in hypoxic cells remained stable over the observed period **Figure 4 B1**). Consistent with this finding, ATP levels in hypoxic cells were only slightly reduced, while ATP levels in normoxic cells decreased with increasing incubation time **Figure 4 B2**). 1,5-(PP)_2_-InsP_4_ and InsP_6_ levels in hypoxic cells were not significantly lower when compared with those in normoxic cells, while over time, a decrease in 1,5-(PP)_2_-InsP_4_ levels was observed under normoxia (**Figure S3 I A1** and **A2**). As the data demonstrate, ATP and PP-InsP levels are dependent on p[O_2_] and also on the duration of the experiment. Such time-dependent adaptations of cellular ATP levels to low O_2_ levels have been observed in earlier studies as for instance by Frezza *et al*. who found decreased ATP levels in hypoxic HCT116 cells (36 h under 1 %O_2_)^[49]^. Moreover, hypoxic adaptation of PP-InsP5 has also been observed in bone marrow-derived mesenchymal stem cells (BM-MSCs): when serum-deprived BM-MSCs were exposed to hypoxia (1 % O_2_), the PP-InsP_5_/IP6 ratio was higher than the PP-InsP_5_/IP_6_ ratio in corresponding normoxic cells.^[50, 51]^ Low p[O_2_] might therefore substantially contribute to alterations of PP-InsP_5_ levels inside serum-deprived hypoxic cells. Moreover, another study discovered increased IP6K gene expression in largemouth bass exposed to hypoxic conditions.^[52]^ Hence, 5-PP-InsP_5_ might be part of the cellular machinery that mediates the metabolic adaptation to reduced p[O_2_].

It is known that cells undergo various metabolic adaptions during hypoxia, for instance upregulation of glycolysis, in order to diminish the respiratory rate. This action prevents ROS overproduction and O_2_ depletion when its levels are limited.^[53]^ In this context, 5-PP-InsP_5_ might have a key role in governing the cellular response to ROS signalling under hypoxia. Decreased cellular ATP utilization under hypoxia^[53]^ explains the stable ATP levels in hypoxic HCT116^UCL^ cells observed over a period of 31 h. In contrast, corresponding normoxic cells maintain their regular energy metabolism likely leading to decreased ATP availability with increasing cell growth (**Figure 4 C**).

Next, the effect of reduced p[O_2_] on PP-InsP concentrations in β-lap treated cells was evaluated. HCT116^UCL^ cells were incubated for 23 h under hypoxia and 10 μM of β-lap was then added to the cells. The cells were incubated with the naphthoquinone for another 2 h under reduced p[O_2_]. Whereas PP-InsP levels in β-lap treated normoxic cells were reduced (ca. 6-fold for 5-PP-InsP_5_, ca. 7-fold for 1,5-(PP)_2_-InsP_4_), PP-InsP levels in β-lap treated cells grown under 1% O_2_ were not changed, clearly highlighting the O_2_ dependence of β-lap action (**Figure 4 C1**). InsP_6_ levels in neither normoxic nor hypoxic HCT116^UCL^ cells subjected to β-lap were affected (**Figure 4 C2**). Moreover, consistent with these findings, cellular ATP levels in hypoxic β-lap treated HCT116^UCL^ cells did not change in response to the drug. In contrast, ATP levels in corresponding normoxic cells subjected to 10 μM of β-lap were decreased (**Figure S3 I B**). These results underscore that β-lap reduces PP-InsP levels *via* ROS production, which is prevented in the absence of O_2_, and not by direct IP6K binding.

### Hypoxia does not influence the effect of other IP6Ks inhibitors

There are a few other known inhibitors of PP-InsP synthesis *in cellulo*, among them the widely-used IP6K inhibitor *N*^2^-(*m*-(trifluoromethyl)benzyl) *N*^6^-(*p*-nitrobenzyl)purine^[46]^ (TNP, structure shown in **Figure S3 II A**) and the natural product Quercetin^[47]^ (Q, structure shown in **Figure S3 II A**), which is an antioxidant.^[54]^ The potency of both, Q and TNP, might also depend on p[O_2_]: Q is able to increase intracellular ROS^[55]^ which might affect the biosynthesis of PP-InsPs and the nitro group of TNP could be reduced under hypoxia.^[56, 57]^ Therefore, we took advantage of the hypoxia chamber to investigate, whether the inhibitory effect of Q and TNP on IP6Ks is dependent on p[O_2_]. Q was capable of significantly reducing 5-PP-InsP_5_ levels under hypoxia as well as normoxia (**Figure S3 II B1 and B2**). Consequently, the redox activity of Q might only marginally contribute to a reduction of cellular PP-InsP synthesis. Regarding TNP, the results demonstrate, that its activity was not affected by reduction of O_2_ levels. Of note, under the applied experimental conditions, TNP reduced PP-InsP levels only to a very minor extent. Therefore, we normalized PP-InsP levels by InsP_6_, which revealed significant but only slight reduction of 5-PP-IP_5_ levels in TNP treated normoxic and hypoxic cells **(Figure S3 II B1-B3**).

### β-Lapachone mediated modulation of PP-InsP levels requires NQO1

Finally, the role of NQO1 in the response of PP-InsP levels to β-lap was investigated. The human breast cancer cell line MDA-MB-231 proved to be useful for this purpose: MDA-MB-231 cells are homozygous for the NQO1*2 polymorphism^[58]^ caused by a C to T point mutation in the NQO1 complementary DNA.^[59, 60]^ Cell types homozygous for the mutant allele have no measurable NQO1 activity and are consequently resistant to β-lap.^[23, 41, 59–61]^ MDA-MB-231 cells were therefore used to study the function of NQO1 in ROS-dependent PP-InsP reduction caused by β-lap. The absence of the NQO1 protein in MDA-MB-231 cells and its presence in HCT116^UCL^ cells were verified by western blot (**Figure 5 A**, whole PAGE shown in **Figure S4**). Also, NQO1-dependent cytotoxicity of β-lap was evaluated (**Figure 5 B1**): cell viability of HCT116^UCL^ significantly decreased when β-lap was applied to the cells in the low micromolar range. In contrast, MDA-MB-231 cells were almost resistant to the naphthoquinone. When dun was tested, comparable results as for β-lap were obtained (**Figure 5 B2**). α-Lap only slightly reduced the viability of HCT116^UCL^ cells and had no cytotoxic effect in MDA-MB-231 cells (**Figure 5 B3**). These findings are in agreement with the literature describing NQO1-dependent lethality for β-lap as well as for dun and minor cytotoxicity for α-lap.^[41,42, 45, 62]^ Also, the cytotoxicity data for the different quinones mirrors their ability to generate intracellular ROS^[42–45]^and to reduce PP-InsP levels (**Figure 3 C1**).

**Figure 5.**
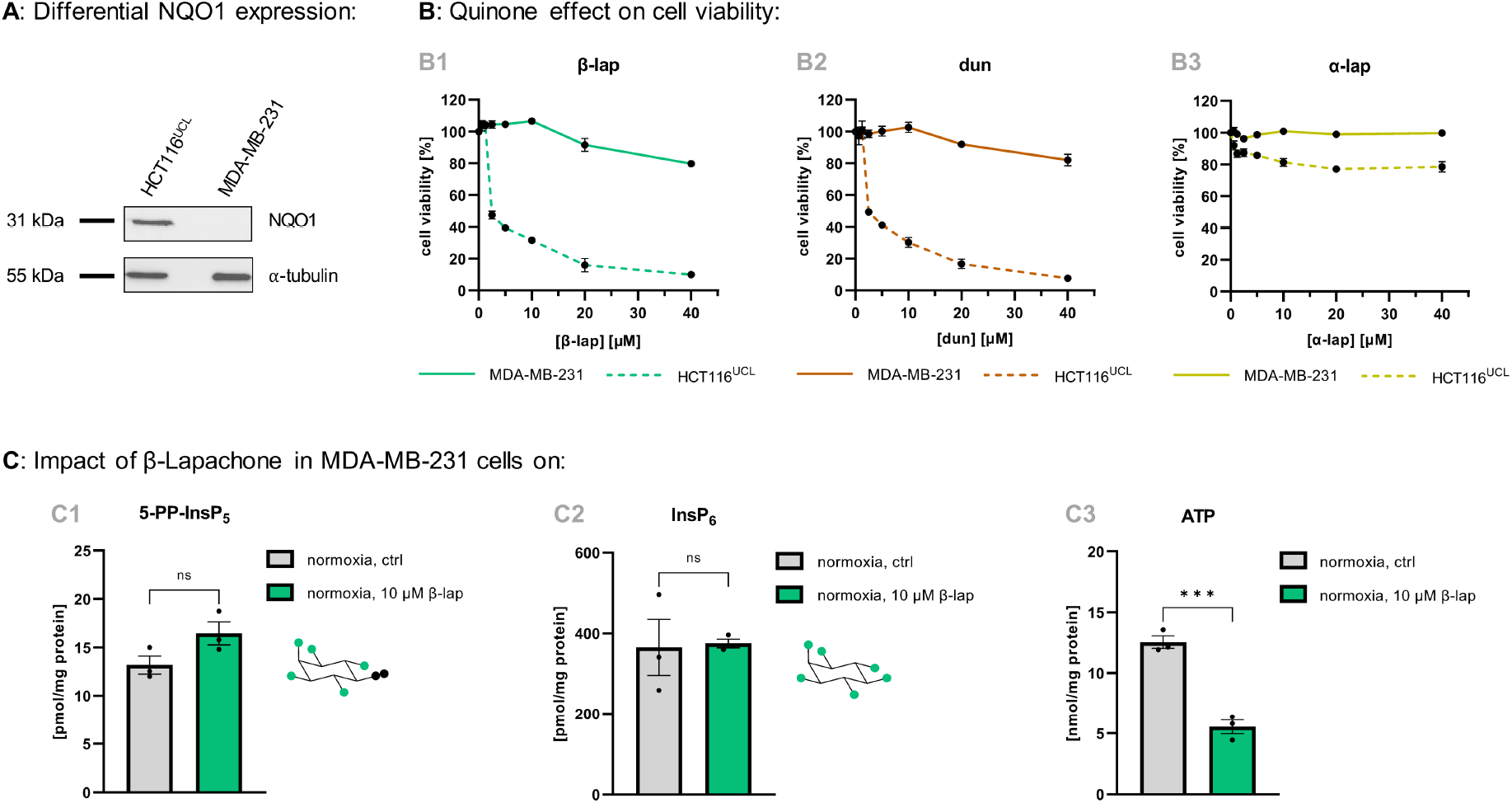
NQO1 is required for β-lap mediated reduction of PP-InsP levels. (***A**) NQO1 protein levels in HCT116^UCL^ and MDA-MB-231 cells*. Equal loading was monitored by using α-tubulin. Whole PAGE is shown in Figure S4 (***B**) Viability assays of MDA-MB-231 cells and HCT116^UCL^ cells treated with different doses of β-lap, dun, and α-lap for 2 h*. After exposure, cells were cultivated for 2 d in compound-free medium before viability was assessed. Data are means ±SEM from three to four replicates. (***C**) 5-PP-InsP_5_, InsP_6_, and ATP levels in normoxic control MDA-MB-231 cells and normoxic MDA-MB-231 cells treated with 10 μM β-lap for 2 h*. ATP was extracted from cells grown in parallel to those used for perchloric acid extration of 5-PP-InsP_5_ and InsP_6_. Data are means ±SEM from three replicates. Statistical analyses to compare treated cells with ctrl cells were performed using an unpaired two-tailed student’s t-test (\*\*\**P* ≤ 0.001). ns: not significant. ctrl: control.

Next, the impact of β-lap on PP-InsPs in NQO1-deficient MDA-MB-231 cells was evaluated. MDA-MB-231 cells were incubated with 10 μM β-lap for 2 h and the cells were then lysed and processed and analyzed as described above. In contrast to the substantially decreased PP-InsP levels in HCT116^UCL^ cells after treatment with 10 μM β-lap (4-8fold, see **Figure 3 A1** and **A2**), 5-PP-InsP_5_ levels in MDA-MB-231 were not affected by the drug, demonstrating again that, *in cellulo*, IP6K is not directly inhibited by β-lap (**Figure 5 C1**). InsP_6_ levels in β-lap treated MDA-MB-231 cells were comparable to those in control cells (**Figure 5 C2**). 1,5-(PP)_2_-InsP_4_ concentrations in this cell line were below the limit of detection. Conversely, ATP levels from MDA-MB-231 cells exposed to β-lap were significantly reduced (**Figure 5 C3**) which had also been observed in HCT116^UCL^ cells subjected to the naphthoquinone (**Figure S2 A**). As MDA-MB-231 cells are not able to express NQO1, loss of ATP levels in β-lap treated MDA-MB-231 cells may be a result of oxidative stress production caused by other one-electron oxidoreductases.^[63, 64]^ Importantly, the significant decrease of ATP in β-lap treated MDA-MD-231 cells had no effect on PP-InsP levels (**Figure 5 C1**). Therefore, the β-lap driven reduction of PP-InsP levels in HCT116^UCL^ is potentially not related to the reduced level of ATP. Instead, it is dependent on a functional NQO1 protein that produces ROS in response to futile cycling of β-lap.

## 5 Conclusion

This study provides insights into how mammalian PP-InsPs are affected by oxidative stress. It was shown here that exposure of NQO1 expressing HCT116^UCL^ cells to exogenous and endogenous ROS reduced PP-InsP levels. Moreover, hypoxia completely abolished β-lap mediated reduction of PP-InsP levels in HCT116^UCL^ cells, which suggests that the quinone is indirectly affecting PP-InsP levels *via* ROS. The loss of potency of β-lap under hypoxia needs to be generally considered when results gained from regular cell culture experiments conducted under normoxia are applied to clinical trials: as physiological O_2_ concentrations typically range from 1 % to 10 %,^[65]^ ROS-dependent β-lap toxicity shown *in cellulo* might be significantly reduced *in vivo*. With regards to the role of β-lap and other modified *o*-quinones as emerging new antitumor agents, we reported that these drugs largely reduced cellular PP-InsP levels while they did not affect InsP_6_. The data obtained from hypoxia experiments showed that basal 5-PP-InsP_5_ and ATP levels in normoxic HCT116^UCL^ cells decreased with increasing incubation time while basal 5-PP-InsP_5_ and ATP levels in co-cultured hypoxic HCT116^UCL^ cells remained stable. This could be explained by the downregulated ATP utilization in hypoxic cells that finally limits ROS overproduction caused by reduced p[O_2_].^[53]^ Hence, 5-PP-InsP_5_ might have a key regulatory function during the cellular adaptation to low O_2_ levels. Further studies are now required to elucidate this hypothesis in more detail. Additionally, it was shown that β-lap did not alter 5-PP-InsP_5_ concentrations inside NQO1-deficient MDA-MB-231 cells. This demonstrates that the modulation of PP-InsPs *via* β-lap requires both, sufficient O_2_ and functioning NQO1 protein. Despite a direct *in vitro* inhibition of IP6K by β-lap, the data provide evidence that, *in vivo*, PP-InsP reductions rely on the NQO1 dependent futile redox cycle of β-lap eventually producing large amounts of ROS.

It remains to be solved how precisely ROS target PP-InsP synthesis on a molecular level. As suggested by Onnebo *et al*. ROS might inactivate the 5-PP-InsP_5_ synthesizing enzymes IP6K1/2/3 *via* the oxidation of cysteine residues inside the protein. In fact, the authors of that study identified a specific evolutionarily conserved cysteine residue that might be attacked by cellular ROS.^[15]^ However, the identification of the specific redox sensitive amino acid *in vivo* will require further investigation.

Among many cytotoxic natural products, β-lap is a promising antitumor agent effective against NQO1-expressing cancer cells. Adding disturbed PP-InsP signalling to its various downstream effects now enables a deeper understanding of its lethality. This offers the potential to design quinone derivatives with increased toxicity and fewer side effects, eventually resulting in more effective and tolerable anti-cancer therapies. Our growing ability to manipulate PP-InsP signalling in conjunction with the use of quinone-based drugs could lead to the development of novel anti-cancer approaches.

## 6 Materials and Methods

### Cell culture

HCT116^UCL^ and MDA-MB-231 cells were cultured in DMEM or DMEM/F12, respectively, each supplemented with 10 % FBS at 37 °C with 5 % CO_2_. Hypoxic experiments were conducted with 1 % O_2_. Details are stated in the SI.

### Assays

Manipulations of PP-InsP levels, cell viability determinations, western blotting, and IP6K *in vitro* experiments are described in the SI.

### CE-ESI-MS analyses

Procedures are based on ref. 29 and described in detail in the SI.

### Statistics

Statistical significance was assessed by unpaired two-tailed student’s t-tests; P ≤ 0.05 is considered significant. For more information refer to the SI.

## 7 Data availability

All data, associated protocols, methods, and sources of materials can be accessed in the text or SI Appendix.

## 8 Acknowledgments

This research was supported by the Deutsche Forschungsgemeinschaft (DFG) under Germany’s Excellence Strategy (CIBSS-EXC-2189-Project ID 390939984). H.J.J. additionally acknowledges financial support from the DFG (Grant JE 572/4-1). HJJ and GL acknowledge funding from the Volkswagen Foundation (VW Momentum Grant 98604). C.L. acknowledges support from the German Scholars Organization and Carl Zeiss Foundation (GSO/CZS 20). The authors wish to thank Dorothea Fiedler (Leibniz Forschungsinstitut für Molekulare Pharmakologie, Berlin) and particularly Minh Nguyen Trung und Robert Harmel for providing ^13^C-labeled reference compounds.

