## supplementary information for "β-Lapachone Regulates Mammalian Inositol Pyrophosphate Levels in an NQO1- and Oxygen-dependent Manner"

**This PDF file includes:**

Supplementary text
Figures S1-S4
SI References

### 1 Abbreviations

| | |
|---|---|
| α-lap | α-lapachone |
| β-lap | β-lapachone |
| BGE | background electrolyte |
| CE | capillary electrophoresis |
| CE-ESI-MS | capillary electrophoresis electrospray ionization mass spectrometry |
| $ddH_2O$ | type 1 ultrapure water |
| DTT | dithiothreitol |
| DMSO | dimethyl sulfoxide |
| DMEM | Dulbecco's modified eagle medium |
| dun | dunnione |
| EDTA | ethylenediaminetetraacetic acid |
| IS | internal standard |
| $InsP_6$ | inositol hexakisphosphate |
| IP6K | inositol hexakisphosphate kinase |
| MRM | multiple reaction monitoring |
| MS | mass spectrometry |
| MTS | 3-(4,5-dimethylthiazol-2-yl)-5-(3-carboxymethoxyphenyl)-2-(4-sulfophenyl)-2H-tetrazolium |
| NQO1 | NAD(P)H:quinone oxidoreductase 1 |
| PBS | phosphate buffered saline |
| PP-InsP | inositol pyrophosphate |
| Q | quercetin |
| Q-TOF | quadrupole time-of-flight |
| rt | room temperature |
| SDS | sodium dodecyl sulfate |
| SIL | stable isotope labeled |
| TNP | $N^2$-(*m*-(trifluoromethyl)benzyl) $N^6$-(*p*-nitrobenzyl)purine |

### 2 Materials and Methods

#### 2.1 General remarks

Commercially available reagents and chemicals were used for all experiments without further purification, unless stated otherwise. Buffers and aqueous solutions were prepared in type 1 ultrapure water (dd$H_2O$). Molecular biology grade reagents and chemicals were used for biological experiments. Stock solutions of the naphthoquinones β-lapachone (β-lap, # L2037 Sigma Aldrich or # HY-13555 MedChemExpress), α-lapachone (α-lap, # BD298310 BLDpharm), dunnione (dun, # BD00939115 BLDpharm), as well as the inositol hexakisphosphate kinase (IP6K) inhibitiors quercetin[1] (Q, # 047268 Fluorochem) and $N^2$-(*m*-(trifluoromethyl)benzyl) $N^6$-(*p*-nitrobenzyl)purine[2] (TNP, # 406170 Sigma-Aldrich) were prepared in dimethyl sulfoxide (DMSO) or ethanol and aliquots were stored at -20 °C. Vacuum evaporation of samples was conducted using a vacuum concentrator (Concentrator plus, Eppendorf).

#### 2.2 *In cellulo* experiments

HCT116 wild type cells from the University College London (further referred to as $HCT116^{UCL}$)[3] were kindly provided by Adolfo Saiardi's lab. MDA-MB-231 cells originating from CLS were acquired from the Signalling Factory (BIOSS Signalhaus, University of Freiburg). $HCT116^{UCL}$ cells were grown in Dulbecco's modified eagle medium (DMEM, # 41966029 Gibco); MDA-MB-231 cells were grown in DMEM/nutrient mixture F12 (# 11320033 Gibco). Medium of both cell lines was supplemented with 10 % fetal bovine serum (# 10270106 Gibco). All stock cell cultures were grown at 37 °C in a 5 % high humidity $CO_2$ atmosphere (referred to as normoxic conditions). Cells were handled under strictly sterile conditions and routinely passed at 1:4-1:20 dilutions using 0.25 % trypsin-ethylenediaminetetraacetic acid (EDTA). Mycoplasma tests were performed quarterly using a mycoplasma detection kit (MycoStrip, InvivoGen); all cell lines were found to be negative. A microplate reader (Tecan Spark 10M) was used for absorbance measurements. Cell viability and cell counts were determined *via* an automatic cell counter (LUNA II, Logos Biosystems). Inositol pyrophosphate (PP-InsP) and inositol hexakisphosphate ($InsP_6$) levels were obtained from cells grown on 150 mm dishes (# 93150 TPP). In order to determine corresponding ATP levels, parallel 60 mm dishes (# 93060 TPP) were prepared. Every biological experiment was performed at least in triplicate. Organic solvents were applied as vehicle during *in cellulo* experiments with naphthoquinones and IP6K inhibitors; their concentrations did not exceed 0.1 % (v/v), except for cell viability assays (final DMSO concentration: 0.2 % [v/v]).

##### 2.2.1 Manipulation of PP-InsP levels *via* exposure to $H_2O_2$ and naphthoquinones

a. Cells for the extraction of PP-InsPs and $InsP_6$ ($HCT116^{UCL}$: 6 mio [2 d experiments] or 5 mio [3 d experiments], MDA-MB-231: 3 mio) were seeded in a 150 mm dish and medium (20-30 mL in total) was added. If desired, cells intended for the extraction of ATP ($HCT116^{UCL}$: 0.85 mio [2 d experiments] or 0.5 mio [3 d experiments], MDA-MB-231: 0.25 mio) were seeded in parallel 60 mm dishes (containing 5 mL medium in total).

b. The cells were incubated under normoxic conditions for 46 h ($H_2O_2$ treatment) or 72 h (naphthoquinone treatment) in total and reached 70-90 % confluence at the end of the experiment. Overall incubation times of 2 d as well as of 3 d were equally suitable for investigation of mammalian PP-InsP, $InsP_6$, and ATP levels.

c. Before harvest, $H_2O_2$ or the naphthoquinone solution was added to the cells. The cells were incubated under normoxic conditions for 30 min ($H_2O_2$) or 2 h (naphthoquinones). The final $H_2O_2$ concentration was 1.0 mM or 0.1 mM and the final naphthoquinone concentration was 2.5 µM, 5 µM, or 10 µM.

d. Cells cultured on 60 mm dishes and intended for ATP extraction were harvested right before processing of cells grown on 150 mm dishes:

- *Harvest of cells grown on 60 mm dishes:* The medium of the cells was removed and the cells were placed on ice. The cells were washed with phosphate buffered saline (PBS) (4 °C) and then scraped in PBS (4 °C). The cell suspension was centrifuged (300 g, 4 min, 4 °C) and the pellet was stored at -80 °C. ATP was extracted following the procedure described in **section 2.2.3**.

- *Lysis of cells grown on 150 mm dishes:* The medium of the cells was removed and the cells were placed on ice. Perchloric acid extraction of PP-InsPs and $InsP_6$ as well as subsequent cell lysis and protein concentration determination was conducted according to the protocol developed by Qiu *et al.*[4] with the following deviations: For cell lysis, 1.5-2.0 mL of lysis buffer (0.1 % sodium dodecyl sulfate [SDS] in 0.1 M NaOH) were used and dishes were incubated on a tilt table until all proteins were solubilized (15-20 min). Isolation products were measured by capillary electrophoresis electrospray ionization mass spectrometry (CE-ESI-MS) as described in **section 2.4**.

##### 2.2.2 Investigation of PP-InsP levels in hypoxic HCT116$^{UCL}$ cells

A HypoxyLab hypoxia workstation (Oxford Optronix) was used for the cultivation of HCT116$^{UCL}$ cells under reduced $O_2$ pressure. For all hypoxic experiments, a gas mixture of 94 % $N_2$, 5 % $CO_2$ and 7 mmHg (~1 % $O_2$) was adjusted and the relative humidity was set to 75 % (referred to as hypoxic conditions). For every experiment, parallel dishes with cells grown under normoxic conditions were prepared. Medium and PBS used for hypoxic cells were pre-equilibrated under hypoxic conditions (at least for 12 h) before usage. Volumes of medium for hypoxia experiments (as well as for parallel experiments under normoxic conditions) were increased (150 mm dish: 35 mL, 60 mm dish: 6 mL) compared to regular *in cellulo* studies under normoxia (**section 2.2.1**). In addition, medium was changed before placing the cells in the hypoxia chamber as hypoxia shifts the metabolism of the cell from oxidative to glycolysis thereby leading to increased lactate production.[5]

a. HCT116$^{UCL}$ cells were seeded in 150 mm dishes and/or 60 mm dishes (for seeding densities refer to **section 2.2.1**). The cells were cultured under normoxic conditions for 22 h. When cells were incubated under hypoxia for a period ≥ 25 , the cell number for 60 mm dishes was decreased to 0.8 mio cells per dish in order to avoid too high confluence at the end of the experiment.

b. After 22 h of preincubation, the medium of the cells intended for hypoxic conditions was removed and the cells were placed in the hypoxia chamber. New pre-equilibrated medium was added to the cells and the cells were incubated under hypoxic conditions for the desired time intervals. Medium of parallel dishes containing cells cultured under normoxic conditions was changed in accordance with the medium of hypoxic cells. Normoxic cells were incubated under normoxic conditions for the same time periods.

c. If desired, the cells were treated with different inhibitors: β-lap (final concentration: 10 µM) was added to the cells 23 h after subjection to hypoxic conditions and the cells were incubated with β-lap for 2 h. Q (final concentration: 2.5 µM) was added to the cells 24 h after subjection to hypoxic conditions and the cells were incubated with Q for 1 h. TNP (final concentration 10 µM) was added to the cells directly after subjection to hypoxic conditions and applied for 48 h.

d. Workup:

*Harvest of cells grown on 60 mm dishes:* Under hypoxic conditions, the medium of the cells was removed, the cells were washed with pre-equilibrated PBS (4 °C) and then scraped in pre-equilibrated PBS (4 °C). The cell suspension was centrifuged (300 g, 4 min, 4 °C) and the pellet was stored at -80 °C. Normoxic cells were processed accordingly on the laboratory bench. ATP was extracted following the procedure described in **section 2.2.3**.

*Lysis of cells grown on 150 mm dishes:* Under hypoxic conditions, the medium of the cells was removed and the cells were washed with pre-equilibrated PBS (4 °C). Perchloric acid (1 mL, 1 M, 4 °C) was added to the cells and distributed equally. The cells were removed from the hypoxia chamber. Alternatively, perchloric acid (1 mL, 1 M, 4 °C) was added directly after removal of the dish from the hypoxia chamber (to avoid reoxygenation before quenching of metabolism). Normoxic cells were treated accordingly on the laboratory bench. The dish was processed as described in **section 2.2.1**.

##### 2.2.3 Methanol extraction of nucleotides

a. The cell pellet was placed on ice and 80 % MeOH/dd$H_2O$ (1 mL, 4 °C) was added. The sample was vortex mixed and then incubated on ice for 10 min.

b. Three freeze-thaw cycles were performed followed by incubation of the sample at -20 °C for 2 h. The sample was then centrifuged (17.000 g, 10 min, 4 °C).

c. The *supernatant* was vacuum evaporated (room temperature [rt]) until the sample was completely dry. dd$H_2O$ (50 µL) was added to the sample, the sample was vortex mixed and then centrifuged (17.000 g, 5 min, 4 °C) to remove residual debris. The supernatant was analyzed *via* CE-ESI-MS as described in **section 2.4**.

d. The *cell pellet* was resuspended in cell lysis buffer (250 µL, 0.1 % SDS in 0.1 M NaOH), vortex mixed and rotated for 15 min at rt. The sample was centrifuged (17.000 g, 5 min, 4 °C) and the supernatant was stored at -80 °C until the protein content was determined via the DC protein assay (Biorad # 5000116) using bovine serum albumin as calibration standard.

##### 2.2.4 Cell viability assays

Cell viability of naphthoquinone treated cells was determined via the 3-(4,5-dimethylthiazol-2-yl)-5-(3-carboxymethoxyphenyl)-2-(4-sulfophenyl)-2H-tetrazolium (MTS) assay (# G5421 Promega).

a. MDA-MB-231 cells or HCT116$^{UCL}$ cells (5000 cells per well) were seeded in a 96-well plate (# 92096 TPP) and medium (100 µL per well in total) was added. The cells were incubated under normoxic conditions for 24 h.

b. Old medium (50 µL) was removed and new medium (50 µL) containing the desired naphthoquinone was added. The final compound concentration ranged from 0.625-40 µM. DMSO was added in analogy to control cells; the final DMSO concentration was 0.2 % (v/v). The cells were incubated under normoxic conditions for 2 h.

c. Compound-containing medium was replaced by compound-free medium (100 µL) and the cells were incubated under normoxic conditions for 43 h.

d. MTS/phenazine methosulfate solution (20 µL) was added and the cells were incubated under normoxic conditions for 3 h in the dark.

e. The absorbance at 490 nm was measured and absorbance values were blank corrected. Cell viability was determined by setting the viability of DMSO treated control cells to 100 %.

##### 2.2.5 Western blotting

a. HCT116$^{UCL}$ or MDA-MB-231 cells (400.000 cells per well) were seeded in 6-well plates (# 657160 Greiner Bio-One) and medium (2 mL per well in total) was added. The cells were incubated under normoxic conditions for 48 h.

a. The cells were harvested *via* trypsinization and the PBS-washed cell pellet was resuspended in radioimmunoprecipitation assay buffer (70 µL, 4 °C; # 39244.01 SERVA) containing protease inhibitor cocktail (10 % [v/v]; # 4693132001 Roche) as well as phosphatase inhibitor cocktails (1 %[v/v] respectively, # P5726 and P0044 Sigma-Aldrich). The cells were incubated on ice for 30 min followed by centrifugation (12.000 g, 10 min, 4°C). The lysate was stored at -20°C until further use.

b. Protein concentrations of whole cell lysates were determined via the BCA assay (# 23227 ThermoFisher) using bovine serum albumin as calibration standard.

c. Cellular proteins (100 µg/µL) in Laemmli buffer (10 % [v/v] glycerol, 2 % [w/v] SDS, 0.01 % [w/v] bromphenol blue, 100 mM dithiothreitol [DTT], 62.5 mM Tris-HCl, pH 6.8) were prepared and the sample was denatured by boiling at 95°C for 5 min. The sample was stored at -20°C until further use.

d. Proteins (15 µg per lane) were separated by a 15 % polyacrylamide gel with a 6 % stacking gel and then transferred to a polyvinylidene fluoride membrane using standard western blotting techniques.

e. Non-specific binding sites on the membrane were blocked and the membrane was incubated with NAD(P)H:quinone oxidoreductase 1 (NQO1) primary antibody (monoclonal mouse, 1:100 dilution; # sc-32793 Santa Cruz Biotechnology) or α-tubulin primary antibody (monoclonal mouse, 1:100 dilution; # sc-32293 Santa Cruz Biotechnology) at 4°C overnight.

f. Detection was performed using IgG horseradish peroxidase-conjugated secondary antibody (anti-mouse, # NA931 Cytiva) at rt for 1 h followed by enhancement with electroluminescence. Secondary antibodies were detected with an Analytic Jena UVP ChemStudio. α-Tubulin was used as loading control to ensure equal loading amounts.

#### 2.3 IP6K1 *in vitro* assays

His-IP6K1 was purified from *E.coli* as described elsewhere[6, 7] and stored in dialysis buffer (100 mM NaCl, 20 mM Tris-HCl, 1 mM $MgCl_2$, 1 mM DTT, 20 % [v/v] glycerol, 0.01 % [w/v] 3-[(3-cholamidopropyl)dimethylammonio]-1-propanesulfonate, pH 7.4). High quality $InsP_6$ is required for IP6K1 reactions and was synthesized in-house as reported previously.[8] Information on the source of stable isotope labeled (SIL) internal [$^{13}$C]-$InsP_6$ and [$^{13}$C]-5-PP-$InsP_5$ standards utilized for *in vitro* assays is given in **section 2.4**. Stock solutions of investigated naphthoquinones in DMSO were diluted with dd$H_2O$ to the desired concentrations before addition to the assay. For *in vitro* reactions, *mixture A* containing 10 × buffer (1 µL; 500 mM NaCl, 200 mM 4-(2-hydroxyethyl)-1-piperazineethanesulfonic acid, 60 mM $MgCl_2$, 10 mM DTT, pH 6.8), purified IP6K1/dialysis buffer (0.75 µL), and dd$H_2O$ (6.25 µL) as well as *mixture B* containing 10 × buffer (1 µL) phosphocreatine (2 µL, 50 mM, # 492378 Fluorochem), creatine phosphokinase (1.5 µL, 400 U/mL, # 2384 Sigma Aldrich),

$InsP_6$ (1 µL, 1 mM), ATP-Mg (2 µL, 10 mM, # A9187 Sigma Aldrich), and $ddH_2O$ (2.5 µL) were prepared on ice.

1. The drug (2 µL, 0.025 mM/0.05 mM/0.1 mM/0.25 mM/0.5 mM in $ddH_2O$) was added to *mixture A* (8 µL) or *mixture A* containing dialysis buffer instead of enzyme (8 µL). Equal volumes of DMSO in $ddH_2O$ were added as vehicle; the DMSO concentration in the final assay volume (20 µL) was 0.25 % (v/v). The assay was mixed by pipetting and incubated for 5 min at rt.
2. *Mixture B* (10 µL) was added, the sample was mixed by pipetting and then incubated for 15 h at 37 °C.
3. The sample was placed on ice and the enzymatic activity was quenched *via* addition of EDTA (2 µL, 100 mM).
4. The sample (5.5 µL) was diluted with $ddH_2O$ (6.5 µL) and spiked with a 1:1 mixture of SIL [$^{13}C$]-$InsP_6$ and [$^{13}C$]-5-PP-$InsP_5$ standards (0.5 µL, 200 µM). The final concentration of the internal standard (IS) in the sample was 8 µM.
5. The sample was analyzed *via* CE-ESI-MS as described in **section 2.4**. The extent of inhibition was calculated as shown in *Equation 1* and *Equation 2*. The respective vehicle control (enzyme + DMSO) was used as reference for each set.

$$5\text{-}PP\text{-}InsP_{5,\ generated} = \frac{[5\text{-}PP\text{-}InsP_5]}{[5\text{-}PP\text{-}InsP_5] + [InsP_6]} \qquad \textit{(Equation 1)}$$

$$inhibition = 1 - \left(\frac{5\text{-}PP\text{-}InsP_{5,\ generated}}{5\text{-}PP\text{-}InsP_{5,\ generated,\ control}}\right) \qquad \textit{(Equation 2)}$$

[5-$PP$-$InsP_5$] = 5-PP-$InsP_5$ concentration inside the sample

[$InsP_6$] = $InsP_6$ concentration inside the sample

6. The extent of inhibition (expressed as percentage) was plotted as a function of the β-lap concentration (expressed in µM) and the $IC_{50}$ was determined with GraphPad Prism software (version 9.2.0) by performing a nonlinear regression based on the variable slope model.

#### 2.4 CE-ESI-MS analyses

CE-ESI-MS analyses of PP-InsPs, $InsP_6$, and ATP from biological samples were conducted based on the procedure as described in the literature.[4, 9] SIL [$^{13}$C]-PP-InsP and [$^{13}$C]-$InsP_6$ standards were provided by Dorothea Fiedler (Leibniz Forschungsinstitut für Molekulare Pharmakologie, Berlin) and synthesized as described elsewhere.[10, 11]

All measurements were conducted on a capillary electrophoresis (CE) Agilent 7100 coupled to a triple quadrupole mass spectrometer (QqQ MS) Agilent 6495c, equipped with an Agilent Jet Stream (AJS) electrospray ionization (ESI) source. A commercial sheath liquid coaxial interface enabled a stable CE-ESI-MS spray, with an isocratic LC pump constantly delivering the sheath liquid.

Capillary zone electrophoresis was performed on a bare fused silica capillary (length: 100 cm, internal diameter: 50 µm) activated with NaOH (1 M, 10 min). Background electrolyte (BGE) was 35 mM ammonium acetate titrated by ammonia solution to pH 9.7. The sheath liquid was composed of a 1:1 dd$H_2O$:isopropanol mixture, with a flow rate of 10 µL/min.

Dried enrichment products containing PP-InsPs and $InsP_6$ were dissolved in dd$H_2O$ (25 µL). PP-InsP and $InsP_6$ extracts (5 µL) and a solution containing SIL IS (5 µL; 4 µM [$^{13}C_6$]-1,5-$(PP)_2$-$InsP_4$, 8 µM [$^{13}C_6$]-5-PP-$InsP_5$, 8 µM [$^{13}C_6$]-1-PP-$InsP_5$, and 40 µM [$^{13}C_6$]-$InsP_6$) were mixed in a CE sample vial. Nucleotide extracts (15 µL) were spiked with a SIL [$^{13}C_{10}$]-ATP IS (0.75 µL, 1 mM). Samples (20 nL) were introduced by applying a pressure of 100 mbar for 10 s. Afterwards, BGE was injected by applying a pressure of 50 mbar for 5 s to receive a BGE plug of 5 nL. A separation voltage of +30 kV was applied, generating a constant CE current of around 19 µA.

The MS source parameters settings were as follows: nebulizer pressure was 8 psi, gas temperature was 150 °C with a flow of 11 L/min, sheath gas temperature was 175 °C with a flow of 8 L/min; the capillary voltage was -2000 V with nozzle voltage 2000 V. Negative high pressure RF and low pressure RF (ion funnel parameters) were 70 V and 40 V, respectively. Parameters for multiple reaction monitoring (MRM) transitions for analytes are shown in **Table S1**.

For determination of PP-InsP, $InsP_6$, and ATP concentrations in biological samples, CE-ESI-MS data was processed using MassHunter Workstation Software (version B.08.00). For graphical representation of electropherograms, OriginPro (version 2018b) was used.

**Table S 1| Parameters for MRM transitions of investigated analytes.**

| compound | precursor ion | product ion | dwell | CE (V) | cell acc (V) | polarity |
|---|---|---|---|---|---|---|
| [$^{13}C_6$]-$(PP)_2$-$InsP_4$ | 411.9 | 362.9 | 30 | 10 | 1 | negative |
| $(PP)_2$-$InsP_4$ | 408.9 | 359.9 | 30 | 10 | 1 | negative |
| [$^{13}C_6$]-PP-$InsP_5$ | 371.9 | 322.9 | 30 | 10 | 3 | negative |
| PP-$InsP_5$ | 368.9 | 319.9 | 30 | 10 | 3 | negative |
| [$^{13}C_6$]-$InsP_6$ | 331.9 | 486.9 | 30 | 46 | 3 | negative |
| $InsP_6$ | 328.9 | 480.9 | 30 | 46 | 3 | negative |
| [$^{13}C_{10}$]-ATP | 516 | 417.9 | 30 | 25 | 1 | negative |
| ATP | 506 | 407.9 | 30 | 25 | 1 | negative |

#### 2.5 Statistical analyses, graphical representations, and drawing software

GraphPad Prism software (version 9.4.1) was used for all statistical analyses and graphical representations of data, unless stated otherwise. Data presents means ±SEM of at least 3 replicates. For statistical analyses, a two-tailed unpaired student's t-test was used (*P ≤ 0.05; **P ≤ 0.01; ***P ≤ 0.001; ****P ≤ 0.0001; ns = not significant).

ChemDraw (version 18.0.0.231) was used for drawing chemical structures and further elements.

### 3 Supporting Figures

Impact of $H_2O_2$ in HCT116[UCL] cells on:

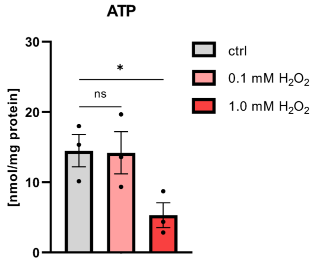

**Figure S1 | Decrease of mammalian ATP levels in response to exogenous ROS ($H_2O_2$).** ATP levels in HCT116[UCL] control cells and HCT116[UCL] cells treated with 0.1 mM or 1.0 mM $H_2O_2$ for 30 min. ATP was extracted from cells grown in parallel to those used for perchloric acid extration of PP-InsPs and $InsP_6$. Data are means ±SEM from three replicates. Statistical analyses to compare treated cells with control cells were performed using an unpaired two-tailed student's t-test (*$P \leq 0.05$). ctrl: control. ns: not significant.

**A**: Impact of β-lapachone in HCT116[UCL] cells on:

**B**: β-Lapachone is effective against IP6K1:

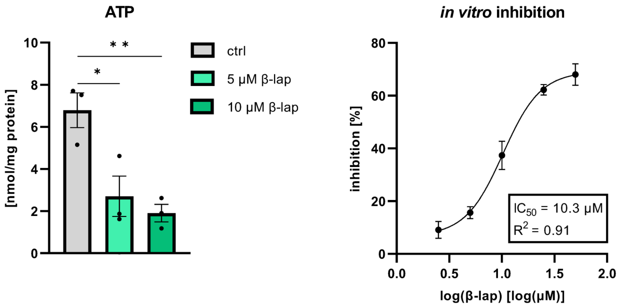

**Figure S2 | Loss of ATP levels in response to β-lapachone generated endogenous ROS and direct IP6K1 inhibition by β-lapachone.** *(**A**) ATP levels in HCT116[UCL] control cells and HCT116[UCL] cells treated with 5 μM and 10 μM β-lap for 2 h.* ATP was extracted from cells grown in parallel to those used for perchloric acid extration of PP-InsPs and $InsP_6$. Data are means ±SEM from three replicates. Statistical analyses to compare treated cells with control cells were performed using an unpaired two-tailed student's t-test (*$P \leq 0.05$; **$P \leq 0.01$). *(**B**) Dose-response curve for the in vitro inhibition of purified IP6K1 via β-lap.* Data are means ± SEM from four replicates. ctrl: control.

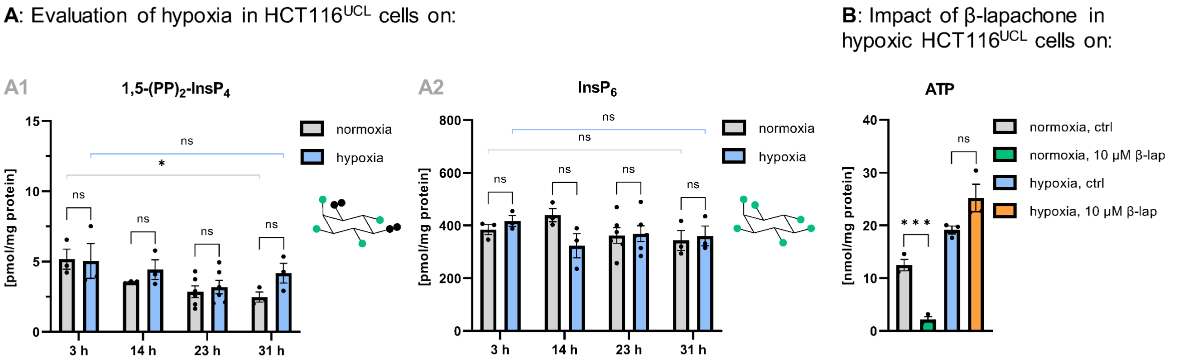

**Figure S3 I | Basal inositol (pyro)phosphate levels under hypoxia and requirement of oxygen for β-lapachone effects on ATP levels.** *(**A**) 1,5-(PP)$_2$-InsP$_4$ and InsP$_6$ levels in HCT116$^{UCL}$ cells grown for 3 h, 14 h, 23 h, and 31 h under normoxic and hypoxic conditions. (**B**) ATP levels in normoxic and hypoxic control HCT116$^{UCL}$ cells and normoxic and hypoxic HCT116$^{UCL}$ cells treated with 10 μM β-lap for 2 h.* ATP was extracted from cells grown in parallel to those used for perchloric acid extration of PP-InsPs and InsP$_6$. Preincubation time of cells under hypoxia prior to drug addition was 23 h. (**A**)+(**B**): Data are means ±SEM from three to six replicates. Statistical analyses to compare hypoxic with normoxic cells, cells cultured for 31 h under normoxia/hypoxia with cells cultured for 3 h under normoxia/hypoxia, and normoxic/hypoxic β-lap treated cells with normoxic/hypoxic control cells were performed using an unpaired two-tailed student's t-test (**P* ≤ 0.05; ****P* ≤ 0.001). ctrl: control. ns: not significant.

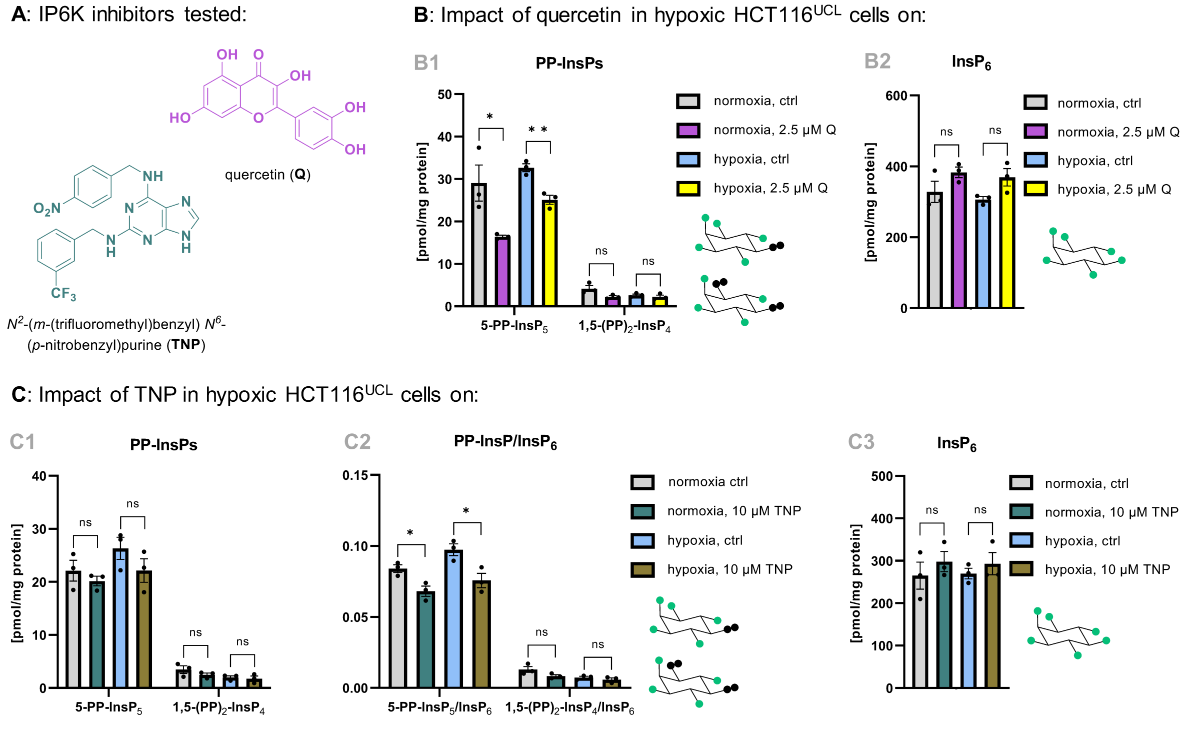

**Figure S3 II | Oxygen independency of mechanistically distinct IP6K inhibitors.** *(**A**) Chemical structures of Q and TNP. (**B**) PP-InsP and InsP$_6$ levels in normoxic and hypoxic control HCT116$^{UCL}$ cells and normoxic and hypoxic HCT116$^{UCL}$ cells treated with 2.5 μM Q for 1 h.* Preincubation time of cells under hypoxia prior to drug addition was 24 h.*(**C**) PP-InsP levels, PP-InsP/InsP$_6$ ratios, and InsP$_6$ levels in normoxic and hypoxic control HCT116$^{UCL}$ cells and normoxic and hypoxic HCT116$^{UCL}$ cells treated with 10 μM TNP for 48 h.* TNP was added at the beginning of the exposure to hypoxic conditions. Data are means ±SEM from three replicates. (**A**)+(**B**): Statistical analyses to compare normoxic/hypoxic inhibitor treated cells with normoxic/hypoxic control cells were performed using an unpaired two-tailed student's t-test (**P* ≤ 0.05; ***P* ≤ 0.01). ctrl: control. ns: not significant.

Full western blot analysis of differential NQO1 expression:

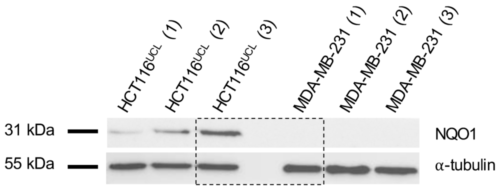

**Figure S4 | Western blot analysis to determine NQO1 protein levels in HCT116$^{UCL}$ and MDA-MB-231 cells.** Three replicates per cell line were analyzed as indicated. Equal loading was monitored by using α-tubulin. The dashed part of the image is shown in **Figure 5 A**.
